# Integrated Genomic Analysis of Hypoxia Genes across Cancer Types Identifies Significant Associations with Cancer Hallmarks

**DOI:** 10.1101/403717

**Authors:** Lingjian Yang, Laura Forker, Christina S. Fjeldbo, Robert G. Bristow, Heidi Lyng, Catharine M. L. West

## Abstract

Hypoxia is a generic micro-environmental factor in most solid tumours. While most published literature focused on *in vitro* or single tumour type investigations, we carried out the first multi-omics pan cancer analysis of hypoxia with the aim of gaining a comprehensive understanding of its implication in tumour biology. A core set of 52 mRNAs were curated based on experimentally validated hypoxia gene sets from multiple cancer types. The 52 mRNAs collectively stratified high- and low-hypoxia tumours from The Cancer Genome Atlas (TCGA) database (9698 primary tumours) in each of the 32 cancer types available. High-hypoxia tumours had high expression of not only mRNA but also protein and microRNA markers of hypoxia. In a pan cancer transcriptomic analysis, ≥70% of the known cancer hallmark pathways were enriched in high-hypoxia tumours, most notably epithelial mesenchymal transition potential, proliferation (G2M checkpoint, E2F targets, MYC targets) and immunology response. In a multi-omics analysis, gene expression-determined high-hypoxia tumours had a higher non-silent mutation rate, DNA damage repair deficiency and leukocyte infiltration. The associations largely remained significant after correcting for confounding factors, showing a profound impact of hypoxia in tumour evolution across cancer types. High-hypoxia tumours determined using the core gene set had a poor prognosis in 16/32 cancer types, with statistical significances remaining in five after adjusting for tumour stage and omics biomarkers. In summary, this first comprehensive *in vivo* map of hypoxia in cancers highlights the importance of this micro-environmental factor in driving tumour progression.

## INTRODUCTION

Hypoxia, low oxygen tension, is a characteristic feature of solid tumours and a hallmark of cancer. Cancer cells adapt to hypoxia via processes including HIF activation and unfolded protein response signalling which alter the expression of genes involved in multiple pathways such as angiogenesis, metabolism, invasion, and epithelial-to-mesenchymal transition ^1,2^. Hypoxia also increases DNA hypermethylation and genome instability ^3,4^. The transcriptional reprogramming generates a pro-survival advantage to hypoxic cancer cells and results in adverse phenotypes, including resistance to therapy and a higher potential to metastasise ^5,6^.

Given its crucial role in tumour progression and resistance to therapy, hypoxia has been explored as a therapeutic target. Approaches for targeting hypoxia have been successfully developed and include increasing oxygen delivery, sensitising hypoxic cells to radiation or using bio-reductive agents that are selectively cytotoxic to hypoxic tumour cells ^7–9^. The effect of hypoxia modification has been supported by high level evidence from clinical trials ^6,10^, making it arguably the most validated target yet to be translated into the clinic ^7^.

Despite increasing efforts to study omics response to hypoxia ^11–15^, most of the literature focused on *in vitro* investigation and limited to single or a small number of tumour types. The *in vivo* impact of hypoxia on tumour genetics and its relationship with other cancer hallmarks are still largely unexplored. To date, the largest *in vivo* multiple cancer types hypoxia gene expression studies were performed by Chi *et al*., ^16^ and Buffa *et al*. ^17^. However, both studies were limited to three cancer types and transcriptome data only. Here we evaluated a set of core hypoxia genes in a large-scale pan-cancer multi-omics analysis. We first defined a core gene set with a high confidence of hypoxia inducibility in multiple cell types. In The Cancer Genome Atlas (TCGA) programme containing mRNA expression of 9698 primary tumours from 32 cancer types, tumours with consistently high expression of core hypoxia genes were identified in each cancer type. High-hypoxia tumours had high expression of other well-known hypoxia markers, including NDRG1 and GAPDH proteins and miR-210 and miR-21 microRNAs. High-hypoxia tumours were highly enriched with the majority of cancer hallmark pathways, including most notably epithelial mesenchymal transition, proliferation and immunology response. By analysing paired clinical and multi-omics data, we also identified significant associations between hypoxia and tumour stage, homologous recombination deficiency, high DNA mutational load and immune cell infiltrate in multiple cancer types. High-hypoxia tumours were associated with a poor prognosis in 16 cancer types. Overall, our work generates the first comprehensive molecular landscape of hypoxia in human cancers, which highlights the important and independent associations with various cancer hallmarks and also profound impact on patient survival.

## RESULTS

### Definition of 52 Core Hypoxia Genes

To curate genes that reliably reflected transcriptional response to hypoxia, we examined hypoxia gene expression experimental studies collated in two recent review papers ^14,15^. The study inclusion criteria were: (1) genome-wide identification of hypoxia responsive genes, and (2) use of multiple cell lines (or tissues) to increase the confidence of finding. Similar studies were combined or filtered out to minimise redundancy. In total ten gene sets ^16,18–28^ were included (more details in Supplementary Methods). Most of the above literature reported genes both induced or suppressed by hypoxia, and we selected only the hypoxia inducible genes, as defined per the original publications. Furthermore, a hallmark hypoxia gene set was obtained from the Molecular Signatures Database ^29,30^, which was derived from four cell line datasets independent from the above ten studies. Another expert curated hypoxia gene set was taken from a review ^1^. The 12 founder gene sets represented experimentally validated transcriptional response genes to hypoxia from nine cancer types, *i.e.* head and neck, cervix, sarcoma, breast, neuroblastoma, glioblastoma, prostate, non-small cell lung cancer, and melanoma. The 12 gene sets also included hypoxia response genes in the following normal cells: B cell, epithelium, astrocytes, monocytes, embryonic cell (Supplementary Table S1). The core hypoxia genes were defined from the founder gene sets as those consistently inducible by hypoxia in multiple (≥*M*) studies. Increasing the value of *M* increases the hypoxia specificity of the core gene set, at the cost of decreasing the coverage of the captured hypoxia transcriptional programme. The 52 genes present in at least four studies (*M*=4) made the final core hypoxia gene set and were used in the subsequent data analysis. Different values of *M* were also attempted with broadly similar results.

The 52 core hypoxia genes overlapped well with most of the 12 founder gene sets (Figure 1A), with the highest overlaps observed with hypoxia genes derived for cervix (79.0%) ^21^, head and neck (74.1%) ^18^, neuroblastoma (68.8%) ^24^ and sarcoma (67.6%) ^20^. Interestingly, all the four gene sets were identified from single cancer types with a large number of cell lines (4 to 11), suggesting that consistently hypoxia inducible genes identified across multiple cell lines in one cancer type are well, albeit not perfectly, preserved. The core hypoxia genes loosely overlapped (12.9%) with the prostate cancer gene set correlating with pimonidazole staining ^23^, which might either reflect a sharply different hypoxia transcriptional programme in profoundly hypoxic prostate carcinoma or a different type of hypoxia measured by pimonidazole ^31,32^. Details of the 52 core hypoxia genes are provided in Supplementary Table S2.

**Figure 1.**
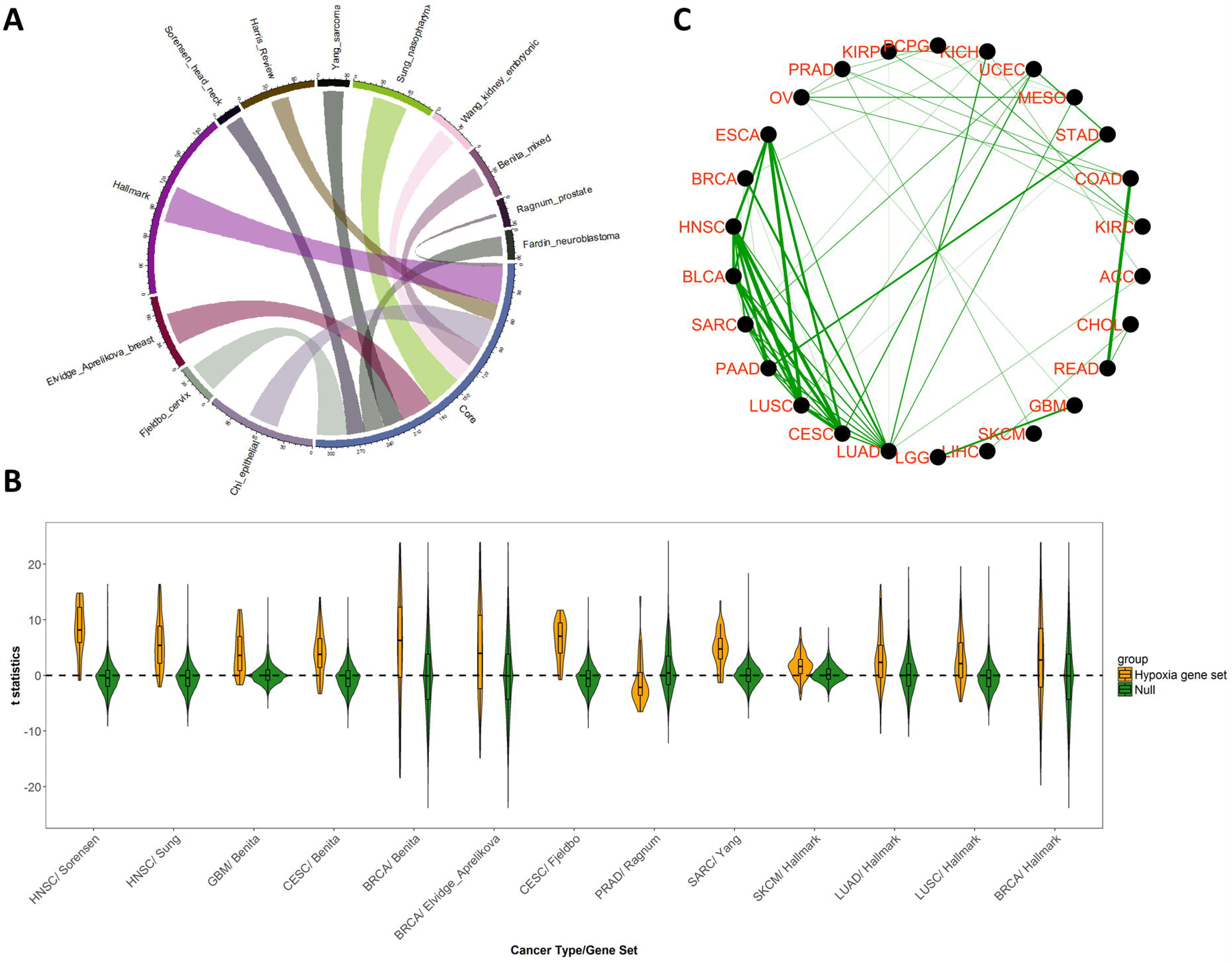
Core hypoxia genes define in vivohypoxia transcriptional programme. **A)** 12 founder gene sets of reliable hypoxia inducible genes were curated from literature. The genes included in more than 4 founder gene sets made up the final core hypoxia gene set. In the Chord diagram, relative circular lengths indicate size of founder gene sets, and thickness of the links between founder gene sets and the core gene set are proportional to the number of member genes from founder set included in the final core gene set. **B)** The distributions of genome-wide *t* statistics from LIMMA analysis comparing high-vs. low-hypoxia tumours were used as null, where the distributions of *t* statistics for 8 founder gene sets were significantly higher than the null distributions. **C)** Similarity of hypoxia transcriptional programme between two cancer types was quantified with Spearman correlation of genome-wide *t* statistics. With an arbitrary cut-off of 0.3, 62 similar pairs could be identified with the strength of association being proportional to the thickness of links.

### Core Hypoxia Genes Identified Tumours with High-hypoxia Phenotype

The expression pattern of the 52 core hypoxia inducible genes were used to collectively define the extent of hypoxia in 9698 human primary cancerous tissues from 32 cancer types in the TCGA, a widely used methodology ^16,17,20,33^. In each cancer type, unsupervised clustering (*K*-means, *K*=2) of the 52 core genes stratified tumours into two groups. In each cancer type, almost all core hypoxia inducible genes were simultaneously elevated in one group of tumours, therefore termed as high-hypoxia gene expression phenotype (Supplementary Figure S1).

We asked whether the high-hypoxia phenotype based on core hypoxia genes captured genome-wide hypoxia transcriptional programme reflected by the founder gene sets in individual cancer types. In each cancer type, genome-wide differential expression analysis between high- and low-hypoxia tumours was performed using LIMMA, where a t statistics was produced for each gene quantifying its relative difference between high-hypoxia and low-hypoxia tumours. Positive *t* statistics indicate up-regulation in high-hypoxia tumours while negative values indicate down-regulation. Among the 12 founder gene sets, eight were derived from samples (cell lines or primary tumours) whose cancer types were among the 32 available in TCGA. For each of those eight gene sets, we compared the distribution of member gene *t* statistics with null distribution of genome-wise *t* statistics in the corresponding cancer type(s). For seven founder gene sets, member genes had significantly higher (t test, P<0.001) *t* statistics compared to the null distributions (Figure 1B), indicating their significant up-regulation in high-hypoxia tumours. Therefore, the results suggest that the identified high-hypoxia phenotype faithfully reflected the transcriptional response to hypoxia in individual cancer types.

We then asked whether an *in vivo* hypoxia transcriptional program was preserved across cancer types. Similarity of the global hypoxia transcriptional program between two cancer types was computed as the Spearman correlation of the genome-wide *t* statistics between high- and low-hypoxia tumours. Using an arbitrary cut-off of 0.3, we identified 62 similar pairs of cancer types (Figure 1C). Our results therefore suggest that molecular responses to hypoxia were conserved across certain cancer types.

### High-hypoxia Phenotype Associated with Cancer Hallmarks

We next sought to identify which biological processes were consistently associated with an *in vivo* high-hypoxia phenotype in a pan cancer setting. Mixed effects linear regression model was applied for each gene, explaining (*Z* standardised) gene expression as a function of hypoxia phenotypes (high- or low-), with cancer type included as a random effect variable. The ~20000 mRNAs were ranked according to their association strengths with hypoxia (coefficients in the regression models), where gene set enrichment analysis (GSEA) ^29^ was employed to identify gene sets whose expressions were enriched or depleted in high-hypoxia tumours.

Among 50 cancer hallmarks, 36 were significantly enriched in high-hypoxia tumours (false discovery rate (FDR)<0.01, Supplementary Table S3), suggesting a large effect of hypoxia on other cancer hallmarks. The top hallmarks enriched in high-hypoxia tumours included not only the well expected hypoxia, glycolysis, angiogenesis and unfolded protein response gene sets, but also epithelial mesenchymal transition (EMT), tumour necrosis factor α (TNF-α) signalling via nuclear factor-αB (NF-κB), cell cycle and proliferation (G2M checkpoint, E2F targets, MYC targets) and immunology response (inflammatory response, interferon gamma/alpha response, allograft rejection, Figure 2). The emergence of EMT as the most enriched process suggests an important role of hypoxia in stimulating tumour invasion and metastasis, consistent with previous literature ^5^. The identified cell cycle and proliferation pathways were recently suggested in the development of high copy number alteration and mutation phenotypes ^34,35^. Our results indicated that some of the genome instability associated with dysregulated cell cycle and proliferation pathways might be due to hypoxia. P53 pathway genes were also significantly up-regulated in the high-hypoxia tumours, in line with a recent report on a p53-dependent pro-apoptotic pathway responding to severe hypoxia ^36^. Moreover, TNF-α and NF-κB signalling control the pro-inflammatory response and have been implicated in tumour cell invasion and metastasis ^37,38^. Our data suggests that hypoxia contributes to tumour progression via activating TNF-α and NF-κB signalling and manipulating inflammatory response. Whether the TNF-α inhibitors could be successfully re-purposed to modify hypoxia in primary cancerous tissues will be of interest ^39^.

**Figure 2.**
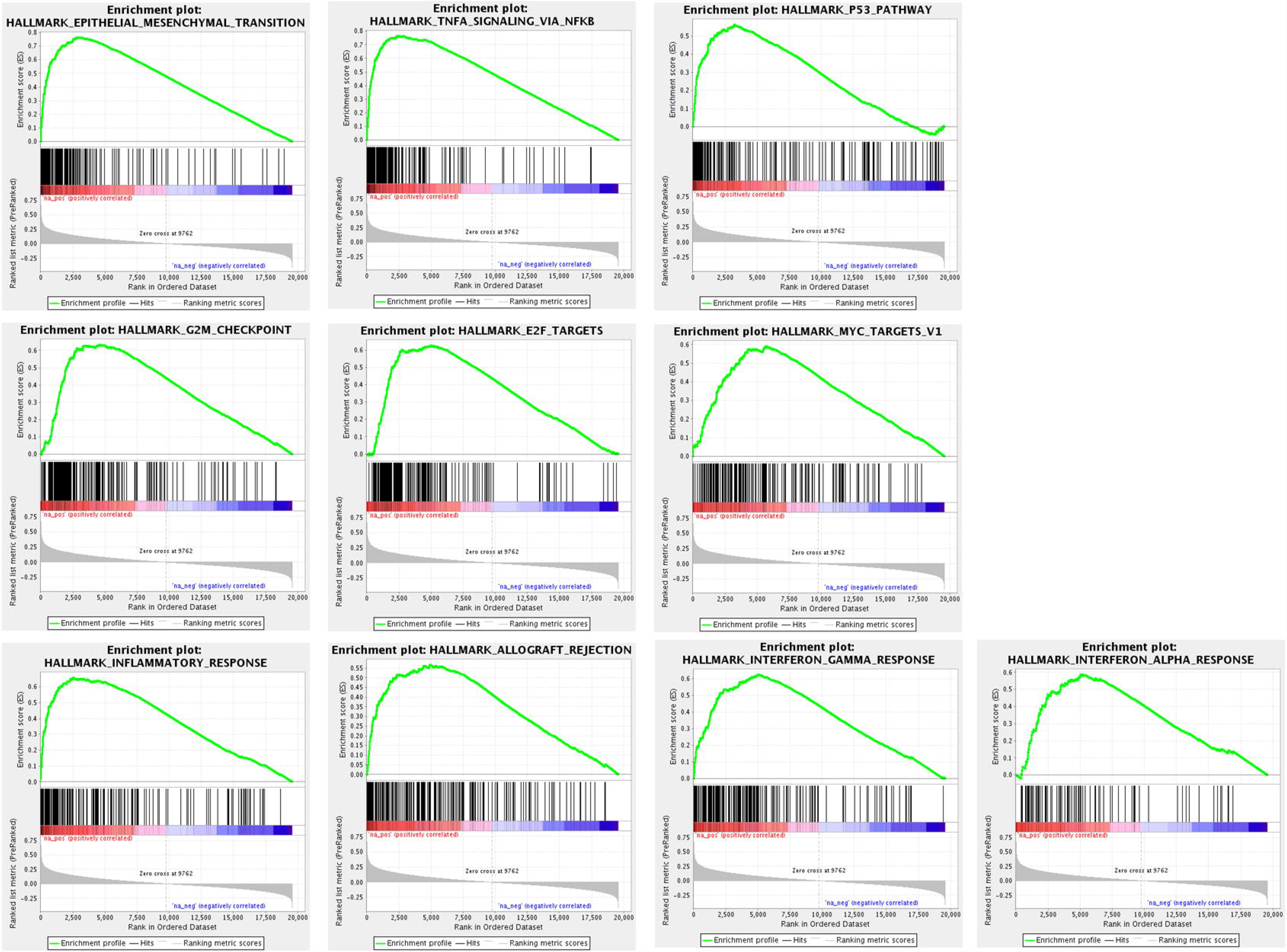
Pan cancer gene set enrichment analysis of the high-hypoxia phenotype. Genes were ranked according to their strength of association with the high-hypoxia phenotype in a pan cancer analysis of 32 cancer types available in TCGA. The high-hypoxia phenotype was defined from expression of 52 core hypoxia inducible genes. GSEA analysis was performed testing the enrichment of 50 hallmark pathways in high-hypoxia tumours. Some of the most enriched pathways, besides hypoxia, glycolysis and angiogenesis, are shown.

As tumour purity might be a confounding factor in the genomic analysis ^40^, we re-ran the above mixed linear regression model with purity corrected estimates. Tumour purity was previously inferred using ABSOLUTE method on copy number alteration data ^34^, and was benchmarked histologically ^41^. Correcting for tumour purity only marginally affected the GSEA analysis results.

### Proteins and MicroRNAs Associated with High-hypoxia Phenotype

To gain more biological understanding of the high-hypoxia phenotype, we analysed functional proteomic and microRNA (miRNA) abundance data for their differential expression between high-hypoxia and low-hypoxia tumours. Abundance of 198 cancer-related proteins and phosphoproteins was measured for TCGA tumours using reverse-phase protein arrays (RPPA) ^42^. To identify the proteins significantly and consistently associated with hypoxia in a pan cancer setting, we again applied a mixed effects linear regression model for each protein, predicting (Z standardised) its abundance as a function of hypoxia phenotype, with cancer type included as a random effect variable. Thanks to the robust statistical framework employed here, we identified 42 and 85 proteins that were respectively up-regulated and down-regulated in high-*vs.* low-hypoxia tumours (FDR<0.01, Supplementary Table S4). The large number of down-regulated proteins in high-hypoxia tumours is consistent with a global decrease of protein synthesis under hypoxia via reduced mRNA translational efficiency ^43^.

Among the top proteins enriched in high-hypoxia tumours, N-Myc Downstream Regulated 1 and Glyceraldehyde-3-Phosphate Dehydrogenase mRNAs (*NDRG1* and *GAPDH*) were included in the core hypoxia gene set. In addition, mRNA of plasminogen activator inhibitor-1 protein (*SERPINE1*) is stress regulated and was shown to be inducible by hypoxia in a murine macrophage cell line ^44^. Accumulation of epidermal growth factor receptor (EGFR) protein under hypoxia and by over-expressing hypoxia-inducible factor 2-α (HIF2-α) was also reported previously in glioma, breast, prostate cancer cell lines ^45,46^. Fibronectin binds to extracellular matrix, and has significant functions in cell adhesion, migration and growth. Hypoxia was shown to induce fibronectin in mouse embryonic stem cells, promoting their proliferation and migration ^47^. Moreover, annexin 1 (ANNEXIN1), involved in anti-inflammatory effects, was found to be hypoxia inducible in a breast cancer cell line and highly expressed in triple-negative tumours ^48^.

Among the proteins most down-regulated in high-hypoxia tumours, B-cell lymphoma 2 (BCL2) is apoptosis-regulating and resides in the mitochondrial membrane. A previous study suggested that hypoxia induced autophagy by up-regulating BCL2 interacting protein 3 like (*BNIP3L*) and BCL2 interacting protein 3 (*BNIP3*) and disrupting the BCL-2-Beclin1 complex, which generates a survival advantage under hypoxia ^49^. Another study showed that down-regulation of BCL2 might be mediated through miR-210, a well-known hypoxia inducible miRNA ^50^. Another protein highly down-regulated in the high-hypoxia gene expression phenotype was estrogen receptor-α (ER-α), which regulates hormone responsiveness and was shown to be inhibited by hypoxia in breast cancer ER-positive cell lines ^51–53^. Our result was also in line with the clinical observation of an anti-correlation between ER-α and HIF-1α ^53,54^. In a phase II trial, HIF-1α was implicated in the resistance to endocrine treatment for ER-α-positive breast cancer patients ^55^. Our results therefore suggest a possibly important association between hypoxia and ER-α in cancer types beyond breast carcinoma, and the potential benefit of adding inhibitors targeting HIF signalling to mediate endocrine sensitivity. Taken together, our pan cancer protein analysis suggests that certain known hypoxia-mediated cellular processes observed previously in individual cancer types could be generalised to multiple cancer types.

MiRNAs are short non-coding RNAs that have important function in mRNA post-transcriptional silencing and are frequently dysregulated in cancers ^56^. Abundance of more than 700 miRNAs were profiled previously for TCGA tumours ^57^. We applied a mixed effects linear regression model to identify the miRNAs consistently differentially expressed between high- and low-hypoxia tumours in a pan cancer setting. We identified 331 miRNAs induced by hypoxia and another 161 miRNAs suppressed by hypoxia (FDR<0.01, Supplementary Table S5). Among the top hypoxia inducible miRNAs identified by the model, hsa-miR-210 is arguably the most well-known hypoxia miRNA biomarker in the literature ^11,12,58^. In addition, hsa-miR-21 was found to be induced by hypoxia in both normal smooth muscle cells and a pancreatic cancer cell line ^59,60^. Also, four of seven miRNAs shown to be hypoxia inducible in two bladder cancer cell lines ^12^ ranked highly in our pan cancer analysis.

Among the miRNAs most suppressed in hypoxia gene expression phenotypes, hsa-miR-139 expression was reported to be lost in invasive breast cancer tumours, while its *in vitro* over-expression suppressed invasion and migration ^61^. Our analysis suggested that the effect of has-miR-139 might be mediated through hypoxia-related signalling. A direct regulatory relationship was also reported between hsa-miR-30a and autophagy-related BCL2 and Beclin-1 in small cell lung cancer and head and neck cancers ^62,63^. Hsa-miR-101 is a well-documented tumour suppressor that is repressed by the pro-inflammatory cytokine IL-1β via the COX2-HIF-1α pathway ^64^. Our analyses thus identify some key miRNAs that are probably universally important in mediating hypoxia responses across cancer types and are potential therapeutic targets.

### High-hypoxia Phenotype Associated Independently with High Tumour Stage, Genome Instability, DNA Damage Repair Deficiency, and Immune Infiltrate

While a role of hypoxia in promoting tumour aggressiveness, metastasis and genome instability was identified from studies on single cancer types, we highlight in this work its importance across cancer types. Our pan-cancer evaluation aimed to quantify associations between a high-hypoxia phenotype and multiple aggressive features characterised in previous studies ^34,65–68^: tumour stage, mutational load, homologous recombination deficiency, leukocyte infiltrate and aneuploidy score. Briefly, mutations were called from whole-exome sequencing data and non-silent mutations per million base pairs were used for mutation rate. Mutation rate was log2 transformed to normalise the skewed distribution. Homologous recombination deficiency score was summarised from three types of homologous recombination deficiency or genome scarring, *i.e.* loss of heterozygosity, large-scale state transitions and the number of telomeric allelic imbalances ^68^. Leukocyte score reflected the total level of tumour immune infiltrate and was derived from methylation data. Aneuploidy score was the number of chromosome arms with arm-level copy number alterations.

American joint committee on cancer pathology stages were used in most cancer types. For CESC, DLBC, OV, UCEC, UCS and THYM, only clinical stages were available and therefore used instead. In eight of the 27 cancer types where tumour stages were available, high-hypoxia tumours were associated with higher tumour stage (Fisher’s exact test, FDR<0.1, Figure 3A). One interesting question is whether hypoxia was associated with higher tumour stage independently of other confounding factors. We performed multiple regression analysis predicting tumour stage (modelled as a discrete variable from 1 to 4) as a function of hypoxia phenotype, mutation load, homologous recombination deficiency score, aneuploidy score and leukocyte infiltrate score. High-hypoxia was independently associated with high tumour stage in nine cancer types (FDR<0.1, Figure 3B). The results, combined with various pre-clinical evidences of hypoxia in increasing tumour aggressiveness ^69^ and the success of hypoxia-targeting treatments in limiting tumour progression ^70^, indicate a role of hypoxia in driving tumour progression across multiple cancer types.

**Figure 3.**
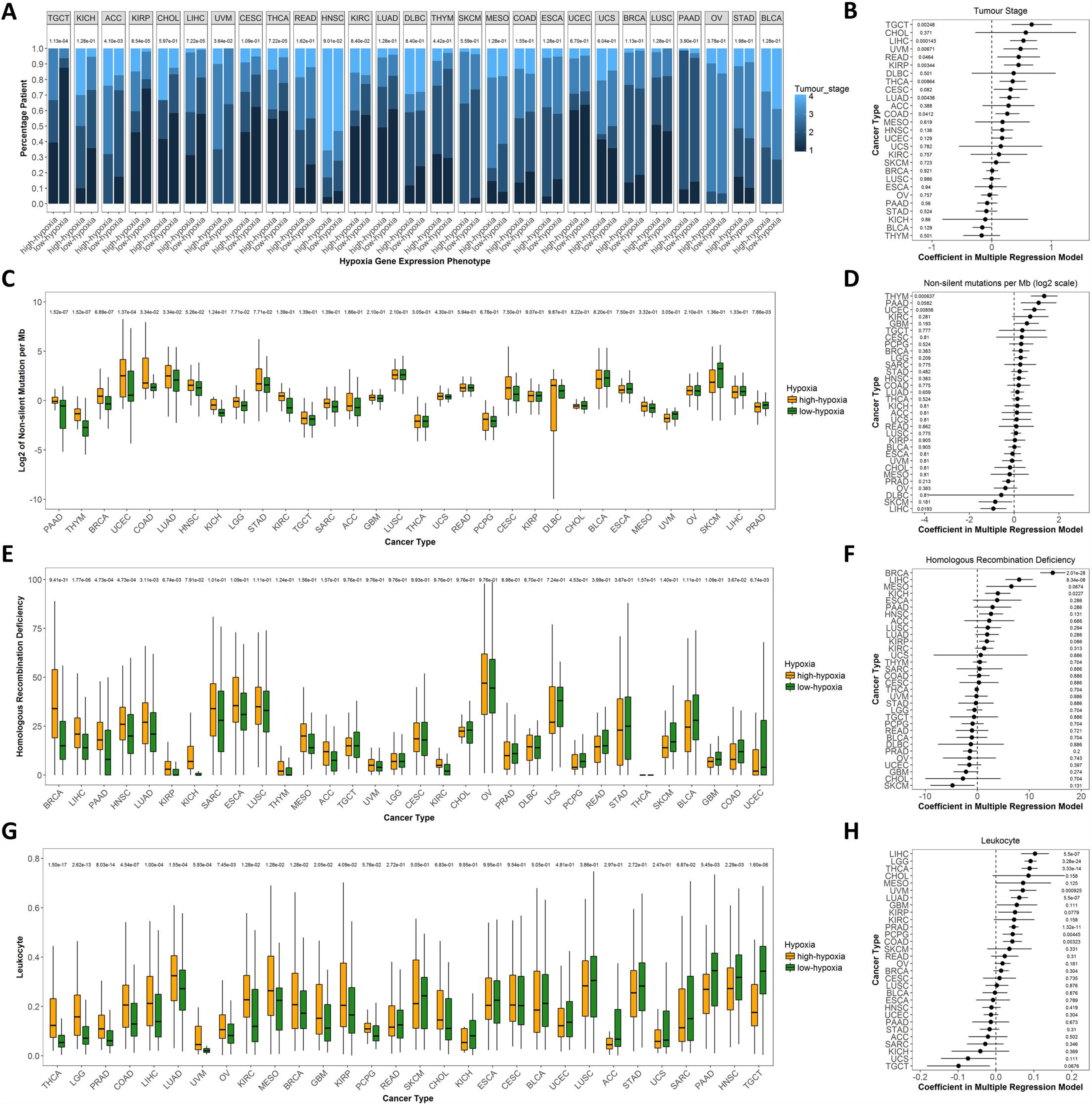
Associations between hypoxia phenotype, tumour stage and other cancer hallmarks in 32 cancer types. **A)** Associations between hypoxia and tumour stages were evaluated using χ^2^ test. **B)** Associations between hypoxia and tumour stages were analysed in multiple regression models adjusting the confounding factors. **C)** and **D)** Associations between hypoxia and mutation rate were analysed using both *t* test and multiple regression models adjusting the confounding factors. **E)** and **F)** Associations between hypoxia and homologous recombination deficiency scores were analysed with both *t* test and multiple regression analyses. **G)** and **H)** Associations between hypoxia and leukocyte were analysed in both *t* test and multiple regression analyses. In each plot cancer types were ranked according to the effect size. Numerical values in each plot indicate FDR values.

Various studies reported increased DNA mutation rate among *in vitro* cell lines exposed to hypoxia ^4^. We observed significantly higher mutation rates in high-hypoxia tumours across nine cancer types (t test, FDR<0.1, Figure 3C). In multiple regression models, hypoxia was adjusted with the above mentioned variables and also tumour stage. Tumour stage was modelled as an ordered categorical variable using stage 1 as the reference level, while the effects of stage 2, 3, and 4 against stage 1 were estimated. The approach was chosen to better model the non-linear associations between tumour stage and molecular biomarkers. Hypoxia was significantly associated with a higher mutation rate independent of other variables in three cancer types, including thymoma, pancreatic adenocarcinoma, uterine corpus endometrial carcinoma (FDR<0.1, Figure 3D).

High-hypoxia phenotype was also associated with increased homologous recombination deficiency in seven cancer types (t test, FDR<0.1, Figure 3E). In multiple regression models, the association with hypoxia was retained in five cancer types (FDR<0.1, Figure 3F).

Immune infiltrate and hypoxia are both important parts of the tumour micro-environment. Our above transcriptome-based hallmark analysis showed associations between hypoxia and various immunology response pathways. To further explore this finding, we evaluated the association between a high-hypoxia gene expression phenotype and leukocyte level determined from methylation data. Overall leukocyte content in the TCGA tumours was estimated using methylation probes which had the greatest difference between leukocyte cells and normal tissues in a mixture model ^66^. High-hypoxia phenotype was associated with significantly more leukocytes in 14 cancer types and significantly less leukocyte levels in four cancer types (t test, FDR<0.1, Figure 3G). After correcting for confounding factors, hypoxia was associated with higher leukocytes in nine (FDR<0.1, Figure 3H) and lower leukocyte levels in one cancer type. While the associations between tumour mutation rate, copy number alteration burden and response to immunotherapy were recently characterised ^34,71^, our data suggested an important yet previously less explored association between hypoxia and the tumour immunological landscape.

All the above analyses were repeated with correction for tumour purity as a confounding factor, which only marginally affected the results (data not shown). Overall, our analyses suggest a very profound association between hypoxia and tumour genetics at multiple molecular levels.

### Hypoxia is A Poor Prognostic factor in Multiple Cancer Types

To investigate the impact of hypoxia on patient survival, we next evaluated the prognostic value of hypoxia gene expression phenotypes using three clinical endpoints of overall survival, disease specific survival and progression free survival ^67^. Patient were treated with surgery, where follow up data were censored at 10-year for each endpoint. In 16 cancer types, hypoxia was a poor prognostic marker in at least one and often multiple clinical endpoints (Cox regression, FDR<0.1 for each endpoint, Figure 4A). Although hypoxia is widely recognised as a strong mediator of resistance to radiotherapy, this analysis emphasises its importance in surgery-treated patients.

**Figure 4.**
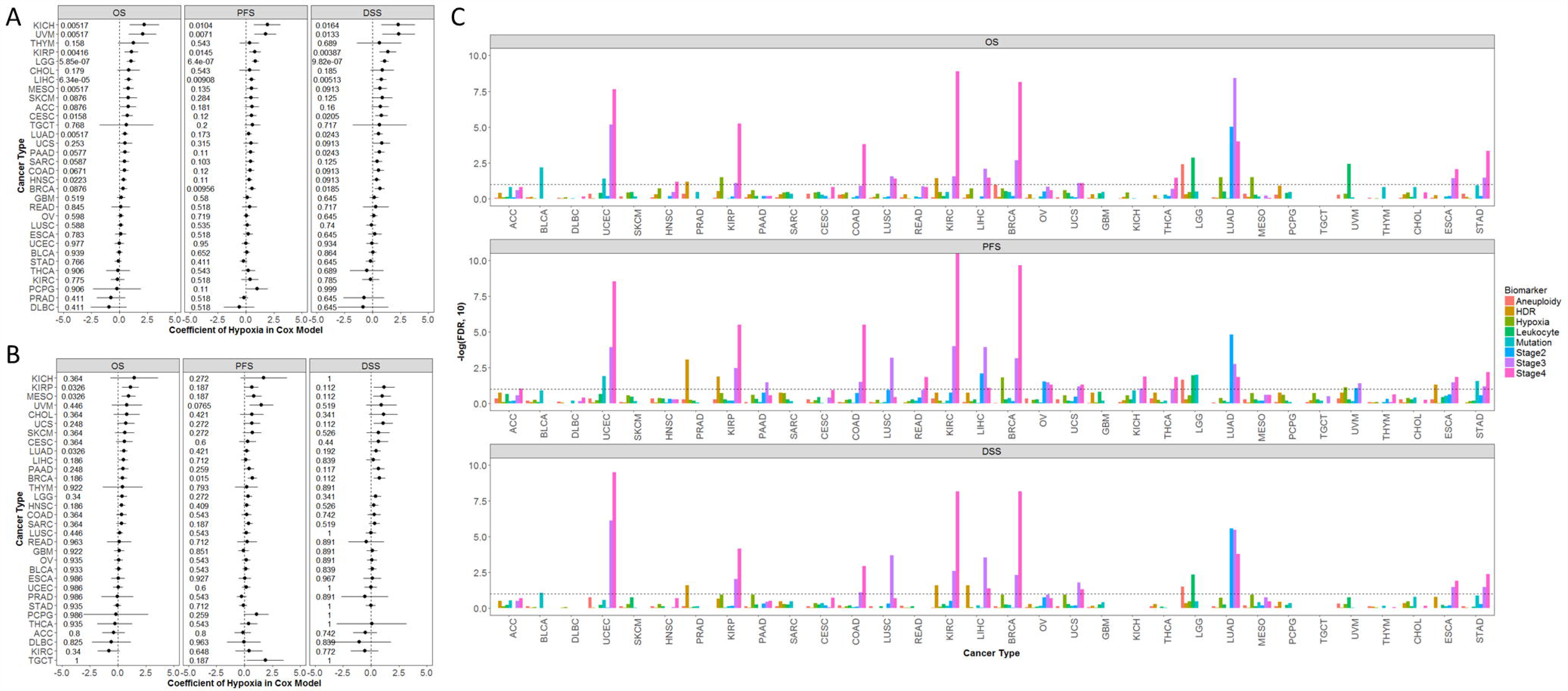
Prognostic value of hypoxia phenotypein 32 cancer types. **A)** Prognostic significances of the 52 core hypoxia gene expression phenotype (high *vs.* low) alone were evaluated in each cancer type using Cox proportional hazard model. The analysis was performed for overall survival, progression free survival and disease specific survival separately. Dots indicate coefficients from Cox model and lines indicate 95% confidence intervals. In each endpoint *P* values were corrected into FDR (values on the side). **B)** The hypoxia gene expression phenotype was adjusted with mutation load, homologous recombination deficiency score, aneuploidy score, leukocyte infiltrate and tumour stage (using stage 1 as reference). **C)** The same multivariable analyses done in **B)**. For each endpoint and each biomarker, *P* values from 32 Cox models were corrected into FDR. Y axis is -log(FDR,10), with higher values indicating more significance. The dashed horizontal line indicates a FDR of 0.1.

We further adjusted hypoxia with mutation load, homologous recombination deficiency score, aneuploidy score, leukocyte infiltrate score and tumour stage in multivariable Cox models. For each endpoint and each biomarker, the nominal *P* values from Cox models were corrected using FDR. In five cancer types (kidney renal papillary cell carcinoma, mesothelioma, lung adenocarcinoma, uveal melanoma and breast invasive carcinoma), hypoxia remained a significant poor prognostic factor for at least one clinical endpoint (FDR<0.1, Figure 4B). Note that in the multivariable analyses, the prognostic significances of a high-hypoxia phenotype were comparable to other omics-based biomarkers (Figure 4C), i.e. high homologous recombination deficiency was associated with poor prognosis in four cancer types; high aneuploidy score was associated with poor prognosis in one cancer type; high mutational load corresponded with a good prognosis in three cancer types and poor prognosis in another; high leukocyte infiltrate was associated with a poor prognosis in two cancer types. Compared with omics biomarkers, tumour stage remained a stronger prognostic factor, in line with a previous report ^72^. Nevertheless, the consistent effect of hypoxia on patient survival again highlights that it is a clinically relevant phenotype worth intervention.

## DISCUSSION

Recent advances in high-throughput technologies and establishment of large consortium like TCGA have enabled integrative and pan cancer analyses of many cancer hallmarks, including somatic mutations ^73^, somatic copy number alterations ^34^, DNA damage repair ^68^ and immunology ^66^. These analyses are generating an increasingly clear picture of the whole spectrum of changes associated with cancer. Understanding the impact of hypoxia on tumour biology is a key object in cancer research. By performing the first multi-omics pan cancer analysis of tumour hypoxia we highlight here the profound association between hypoxia and various cancer hallmarks.

It is well accepted in the literature that hypoxia results in large-scale change in transcriptional reprogramming, and the changes differed across cancer types ^15,16,18^. By surveying a large number of experimentally validated hypoxia gene sets, we identified a group of core genes consistently inducible by hypoxia. Note here that we chose to exclude hypoxia gene signatures, as their bioinformatics derivation methods differ substantially and could well reflect other biological parameters, as discussed in a previous review ^14^. We showed that the primary tumours stratified as high-hypoxia by the highly conserved core gene set not only highly expressed the founder cancer-type-specific hypoxia gene sets in most of the cancer types (albeit perhaps not prostate carcinoma), but also were highly enriched with well-known protein and miRNA hypoxia markers. The results therefore showed that we have identified primary tumours with differential levels of hypoxia. Our pan cancer analysis also suggested that some of the hypoxia-mediated mechanisms reported in individual cancer types or *in vitro* could be generalised to multiple cancer types and *in vivo*.

We observed remarkable associations between hypoxia and many other cancer hallmarks, including high mutation rate and more DNA damage repair deficiency. Our data also showed that high-hypoxia tumours had high epithelial mesenchymal transition potential and cell proliferation signatures, consistent with the literature evidence from single tumour sites ^46,74–76^. Previous studies characterised the relationship between somatic mutations and copy number alterations with an immune landscape ^34,71^. Our analysis showed hypoxia was mostly associated with increased leukocyte content, independent from other confounding factors. It is of interest for future research to investigate whether hypoxia biomarkers could improve the prediction of response to immunology therapy.

Hypoxia has been well recognised as mediating resistance to radiotherapy ^36,50,77^ and is arguably the most established target not yet translated into the clinic ^7^. Although studies in some cancer types showed hypoxia was an adverse prognostic feature following surgery ^11,58,78,79^, we showed here its widespread importance for surgery-treated patients across tumour types. Importantly, the effect of hypoxia on surgical outcomes was comparable with other omics marker. Targeting hypoxia and hypoxia signalling, either alone or in combination with other therapies, could well generate a sizable survival benefit for cancer patients.

In summary, our study highlights a high level of conservation of hypoxia response genes across tumour types. Hypoxia has a large effect on multiple cancer hallmarks with EMT, proliferation and immunology response being the most consistently enriched phenotypes. High-hypoxia tumours also have high tumour stage, mutational burden, homologous recombination deficiency and immune cell infiltrate. Our work supports the importance of translating hypoxia modification therapies into the clinic to improve patient survival rates.

## METHODS

### TCGA Omics Data

TCGA is a large-scale cohort for multi-omics analysis of tumours. Recently, TCGA has published the pan-cancer atlas, where all omics data were harmonised for consistent quality control, batch effect correction, normalisation, mutation calling, and curation of survival data. For this work, we extracted processed RNA sequencing V2, microRNA sequencing, reverse phase protein array and patient clinical and survival data from the Pan-cancer Atlas consortium, which were described elsewhere ^34,66–68^. Only the primary tumours were kept for analysis. We also downloaded paired mutational load, homologous recombination deficiency, aneuploidy score, leukocyte infiltrate, tumour stage and patient survival data from the consortium for evaluating the association with hypoxia.

### Stratification and Analyses of Hypoxia Gene Expression Phenotypes

For each cancer type, primary tumours were stratified into two groups (high- and low-hypoxia) by applying unsupervised *K*-means clustering (*K*=2) on the mRNA expressions of the core hypoxia genes. We used kmeans function (1000 random starts) in R (v3.4.1). Differential expression analysis was performed using LIMMA (v3.34.9). To identify the genes consistently associated with high- and low-hypoxia tumours in pan cancer setting, a mixed effect linear model was applied for each gene: *Expression = β* * *Hypoxia* + (1|*CancerType*), where hypoxia (1 indicates high-hypoxia and 0 low-hypoxia status) is a fixed effect variable and cancer type a random effect variable. nlme library (v3.1-131) in R was used for estimating β coefficients, which reflect the sign and strength of associations between gene expression and hypoxia. Expressions of each gene were *Z* standardised to ensure β coefficients were comparable between different genes. When correcting tumour purity as confounding factor, purity value was added as a fixed effect variable: *Expression* = *β*\* *Hypoxia* +*γ* * *Purity* + (1|*Cancer_type*). Genes were ranked from high to low based on β coefficients. GSEA analysis was performed using gene pattern public server and with 50 hallmark gene sets.

### Statistical Analysis

Unequal variance *t* test was used to compare if the mean of two distributions were equal. Fisher’s exact test was used to test independence of distribution of high- and low-hypoxia tumours across different tumour stages. Survival estimates were performed using the Kaplan-Meier method, and hazard ratios (HR) and 95% confidence interval (CI) were obtained using the Cox proportional hazard model. All *P*-values were two-sided and multiple testing corrections were done using false discovery rate. All statistical analyses were done in R (v3.4.1).

